# Direct visualization of transcriptional activation by physical enhancer–promoter proximity

**DOI:** 10.1101/099523

**Authors:** Hongtao Chen, Miki Fujioka, James B. Jaynes, Thomas Gregor

## Abstract

A long-standing question in metazoan gene regulation is how remote enhancers communicate with their target promoters over long distances. Combining genome editing and quantitative live imaging we simultaneously visualize physical enhancer–promoter communication and transcription in *Drosophila* embryos. Enhancers regulating pair rule stripes of *even-skipped* expression activate transcription of a reporter gene over a distance of 150 kb. We show in individual cells that activation only occurs after the enhancer comes into close proximity with its regulatory target and that upon dissociation transcription ceases almost immediately. We further observe distinct topological conformations of the *eve* locus, depending on the spatial identity of the activating stripe enhancer. In addition, long-range activation results in transcriptional competition at the endogenous *eve* locus, causing corresponding developmental defects. Overall, we demonstrate that sustained physical proximity and enhancer–promoter engagement are required for enhancer action, and we provide a path to probe the implications of long-range regulation on cellular fates.

## I. INTRODUCTION

Enhancers play a key role in the control of gene expression and development (Benoist and Chambon, 1981; Levine, 2010; Long et al., 2016). These 50–1500 base pair cis-regulatory elements stimulate transcription from core promoters in a time and tissue specific manner by recruiting context-dependent transcriptional activators and repressors (Levine, 2010; Buecker and Wysocka, 2012). Functions of enhancers are thought to be independent of their position and orientation. Enhancers can act in cis and in trans (Lee and Wu, 2006; Markenscoff-Papadimitriou et al., 2014), can be located upstream or downstream of the core promoter as well as in introns, and as far as a Mb from the promoter (Long et al., 2016). Whole-genome methods have shown that the human genome is riddled with enhancers, with estimates ranging from 200,000 to over a million (Consortium, 2012). Importantly, a significant fraction of these enhancers are located at large genomic distances from the promoters they regulate (Tolhuis et al., 2002; Uslu et al., 2014; Zhang et al., 2013). Even for a compact genome like *Drosophila melanogaster*, at least 30% of enhancer–promoter interactions occur over 20 kb, and in many cases over intervening genes (Arnold et al., 2013; Ghavi-Helm et al., 2014; Kvon et al., 2014).

In this paper we focus on the mechanism by which enhancers communicate with their target genes over large distances. Several models have been proposed (Benabdallah and Bickmore, 2015; Blackwood and Kadonaga, 1998), the most popular of which involves looping of the flexible chromatin polymer such that distal enhancers come into direct contact with their promoter targets. Early evidence from genetic experiments has already revealed the presence of functional long-range enhancer–promoter communication in both vertebrates and flies (Lettice et al., 2003; Lewis, 1978; Qian et al., 1993; Sagai et al., 2005). Recently, chromatin conformation capture (3C) based approaches, involving the cross-linking of chromatin, revealed the physical proximity of distal enhancers and promoters in a variety of vertebrate cells and tissues (Kagey et al., 2010; Mifsud et al., 2015; Spitz, 2016). Yet, despite these demonstrations of both functional and physical interactions, fundamental questions about how enhancers communicate with their target promoters over large distances and how they activate transcription still remain (Levine et al., 2014).

A major gap in our understanding pertains to the role of physical enhancer–promoter interactions in transcriptional regulation. On the one hand, 3C-based experiments have revealed extensive enhancer–promoter interactions that are conserved among developmental stages, cell fates or evolution, suggesting a permissive role of the physical enhancer–promoter interactions on transcriptional activity (Ghavi-Helm et al., 2014; Jin et al., 2013; Sanyal et al., 2012). On the other hand, lineage-specific enhancer–promoter interactions are found to be prevalent in many developmental contexts, arguing for the possibility of an instructive role (Dixon et al., 2015; Javierre et al., 2016). Part of the reason for this discrepancy lies in the fact that the static picture gained from currently employed techniques can only provide correlative evidence. Furthermore, most genomic experiments rely on bulk assays, which might cover temporal and spatial heterogeneity within the samples.

Crucially, a direct dynamic link between enhancer–promoter proximity and transcriptional activation is lacking. Is proximity needed for transcriptional activation, or is it a consequence of transcriptional activity? Even if the most popular hypothesis is that proximity is necessary for transcriptional activation, the above discussed studies either measure transcriptional activity or physical proximity, and experiments that measure both transcription and physical proximity simultaneously have not been reported. What degree of physical proximity is needed for activation? Is there a topological distinction at the gene locus when different enhancers are activating the gene? It is further unclear whether physical proximity is needed transiently for a limited time interval to establish an active state—a state that can be remembered once proximity is lost—or whether continued transcription requires that the enhancer and promoter remain in sustained proximity. To address these fundamental questions an approach that simultaneously captures the dynamics of enhancer–promoter distances and the dynamics of transcriptional activity is necessary.

Here we have devised an assay that uses a combination of genome editing, genetics, and live single-cell imaging to visualize the relationship between enhancer activation of transcription and physical proximity in real time. We show that activation of transcription can only occur when the enhancer comes into close proximity with its regulatory target and that transcription ceases almost immediately when they move apart. Moreover, we find that proximity alone is not sufficient for enhancer action; rather enhancer action requires a direct engagement of the enhancer with its target promoter. Our results suggest that establishment of physical interactions between enhancers and promoters can be the key rate-limiting step in gene regulation over long distances, and we can exclude mechanisms of transient enhancer–promoter associations leading to stable transcriptional activity.

## II. RESULTS

### Genetic Design: *Homie*-dependent long-distance regulation

To determine whether physical proximity is central to enhancer–promoter communication we have taken advantage of a characteristic property of boundary (insulator) elements in flies, namely their ability to pair with themselves, often over large genomic distances. For this purpose we selected a boundary called, *homie*, which marks the 3 end of the *even-skipped (eve)* locus (Fujioka et al., 2013; Fujioka et al., 2009). Genetic studies have shown that *homie–homie* self-pairing interactions can orchestrate enhancer activation of a reporter at distances of at least 2 Mb (Fujioka et al., 2013).

In our experimental system a transgene consisting of the *eve* promoter (no enhancers) and the *lacZ* coding sequence is inserted at an attP site located 142 kb upstream of the *eve* gene (Fig. 1A). When *homie* is included in this reporter transgene, physical interactions between it and the *homie* boundary at the endogenous *eve* locus can be detected in chromosome conformation capture experiments (Fujioka et al., 2016). In fixed embryos sporadic expression of *lacZ* mRNA is observed solely within the limits of the endogenous *eve* stripes (Fig. 1B and 1C). These findings indicate that the activation of the *lacZ* reporter depends on the enhancers in the *eve* locus 142 kb away, and that at this stage in development, the promoter of the reporter has no spontaneous activity nor does it respond to enhancers near the site of insertion.

**Fig 1:**
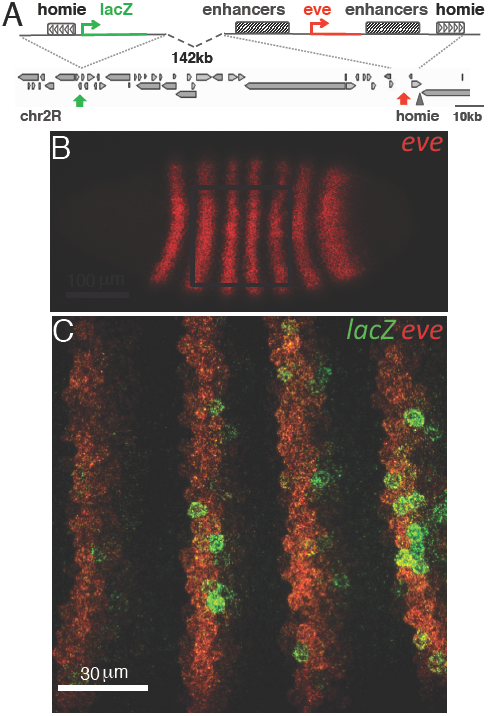
An endogenous genomic construct is designed to investigate long-distance enhancer–promoter interactions. **(A)** Scheme of the construct design and its genomic context. Ectopic *homie* insulator sequence with an *eve* promoter driving *lacZ* is integrated at −142 kb upstream of the endogenous *eve* locus in the Drosophila genome. **(B)** Upper panel: surface view of a 2.5 h old Drosophila embryo hybridized with red fluorescent *eve* mRNA probes. Lower panel: z-stack projection of the marked region in the upper panel. LacZ activity (revealed using green fluorescent *lacZ* mRNA hybridization probes) only occurs sporadically within the limits of the *eve* pattern (red).

### Visualization of transcription and enhancer–promoter dynamics

Key to exploring the connection between enhancer action and physical enhancer–promoter proximity is being able to simultaneously visualize the location of the promoter, the location of the enhancers, and the transcriptional activity in living embryos. For this purpose we introduced tags into the endogenous *eve* gene and the *lacZ* reporter transgene at −142 kb (Fig. 1A). First, we used genome editing to insert an MS2 stem loop cassette (Bertrand et al., 1998; Larson et al., 2011; Yunger et al., 2010) into the 1st intron of the *eve* gene. Maternally expressed MS2 coat protein (MCP) fused to a blue fluorescent protein was used to visualize not only nascent *eve* transcripts but also mark the nuclear location of the *eve* gene and the associated *eve* enhancers. Second, to visualize transcriptional activity of the *lacZ* reporter we placed a PP7 stem loop cassette (Fukaya et al., 2016; Hocine et al., 2013) near the 5 end of the *lacZ* coding sequence. Nascent transcripts expressed from the reporter can then be visualized by the binding of maternally expressed PP7 coat proteins (PCP) fused to a red fluorescent protein (Fig. 2A). Finally, to mark the position of the *lacZ* reporter independent of whether the reporter is active, we used the recently developed parS/parB DNA labeling system (Dubarry et al., 2006; Saad et al., 2014). This system employs *Burkholderia* parS DNA sequences that nucleate the binding of a ParB-GFP fusion proteins, and it is thought to be less disruptive to the local chromatin structure than more traditional DNA labeling systems (Bystricky, 2015).

**Fig 2:**
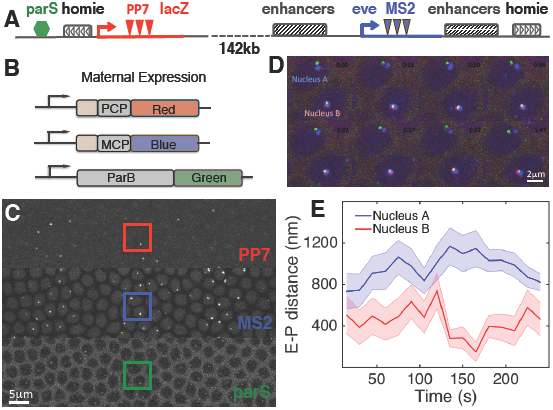
Visualization of enhancer–promoter movements producing transcriptional activity using three-color live imaging. **(A)** Editing the endogenous *eve* locus. 24xMS2 stem loops are knocked into the 1st intron of *eve* to measure *eve* transcriptional activity as well as the position of the locus. 24xPP7 stem loops are inserted into the *homie-lacZ* transgene to monitor *lacZ* transcription, and a parS tag is introduced next to the ectopic *homie* sequence for monitoring the movement of the reporter locus at −142 kb. **(B)** Design of genetic constructs: PCP::red, MCP::blue, and ParB::green fusion proteins that bind PP7 and MS2 stem loops, and *parS* sequences, respectively, are provided maternally. **(C)** Snapshot of an F1 embryo from females carrying all fluorescent proteins in (B), mated with males carrying the genomic construct shown in (A). The embryo displays fluorescent foci for MS2 (blue), PP7 (red) and parS (green) in the corresponding channels. **(D)** 8 snapshots of a time course following two nuclei for 2 min. Nuclei A and B have *eve* activity (blue); nucleus B also has *lacZ* activity (red). **(E)** Instantaneous enhancer–promoter (E-P) distance between endogenous *eve* enhancers (as measured by blue signal) and the transgenic reporter (as measured by green signal) as a function of time for nuclei A (blue) and B (red) from (D). Error bar corresponds to measurement error. See also Fig. S1 and Fig. S2.

Using three-color time-lapse confocal microscopy we captured stacks of optical sections of the surface of two-hours old embryos carrying the tagged *eve* locus and the *parS-homie-lacZ* reporter. In these stacks we can clearly identify individual fluorescent foci in 70-100 nuclei simultaneously (Fig. 2B). In the blue channel we observed the endogenous activity and transcriptional dynamics of the *eve* gene in its characteristic seven-striped pattern. This pattern is quantitatively identical to that observed from the endogenous *eve* gene (Fig. S1). In the green channel we saw the parB foci in all of the nuclei in the embryo, which trace the movement of the *lacZ* reporter within the nucleus. Finally, in the red channel we observed *lacZ* expression in a subset of nuclei in the (blue) *eve* stripes. Consistent with the results from our fixed embryos, *lacZ* expression is restricted to nuclei that reside within one of the seven *eve* stripes.

To test our ability to reliably and accurately measure the localization of the reporter and the *eve* gene we generated a synthetic construct (localization control) in which all three fluorescent proteins are localized within a genomic distance of 2.0 kb (Fig. S2A). By analyzing embryos carrying this construct, we were able to calibrate chromatic aberrations originating from the microscope and also estimate errors in measuring spot localization (Fig. S2B-H). We also tested whether our method of marking the location of the reporter introduces perturbations in the chromatin structure that would hinder our analysis. For this purpose we placed the *parS* sequence at different locations relative to the *lacZ* reporter (Fig. S3A). We also implemented the more traditional *lacO* /LacI system to mark the location of the reporter in the nucleus (Gasser, 2002; Sinclair et al., 2010). No significant difference in chromatin dynamics and transcription kinetics were observed when the *parS* tag was placed at different locations or replaced by the lacO tag (Fig. S3B-H). Based on these analyses, we conclude that our genomic labeling approach allows us to measure chromatin dynamics with an error of 180±6 nm (mean±SEM).

Our initial visualization of reporter activity revealed a close connection between transcription and physical proximity. In nuclei in which the reporter is inactive, the reporter is well separated from the *eve* gene. In this case the green focus, which marks the *parS* sequence in the reporter, and the blue focus, which tags the transcriptionally active *eve* gene are far from each other, and since there is no red focus the reporter is silent (Fig. 2C). In nuclei in which the *lacZ* reporter is ON, the *parS-homie-lacZ* transgene co-localizes with the *eve* gene. In these nuclei the blue, green and red fluorescent foci appear to be attached together (Fig. 2C). This difference is recovered in measurements of the instantaneous spatial distance between the blue and green foci. In nuclei where red reporter activity is present, the distance between the blue and green foci is significantly shorter than the blue-green distance in nuclei lacking the red focus throughout the interval of observation (Fig. 2D and Fig. S3B-C).

### Spatial proximity is necessary for enhancer action

To provide a more detailed picture of how enhancer action is related to spatial proximity, we measured the physical separation between the *eve* gene (blue) and the *lacZ* reporter (green) in embryos carrying the experimental transgene, *parS-homie-lacZ*. We analyzed live images of 2528 nuclei across 35 individual embryos over a 30 min period in nuclear cycle 14 (Fig. 3A). We observed a bi-modal distribution for the time-averaged physical distance (root-mean-squared (RMS) distance, see Methods) that could be modeled as a mixture of two Gaussians (Fig. 3B) at 743±96 nm and at 361±75 nm, respectively (mean±STD).

To confirm that the linkage of the reporter to *eve* is dependent on *homie* we replaced the *homie* sequence with lambda DNA of the same length. We generated a *parS-lambda-lacZ* transgene and analyzed movies of 870 nuclei across 12 individual embryos, again for 30 min in nuclear cycle 14 (Fig. 3A, yellow). For this data set, the distribution of the MS2-parS (blue-green) RMS distance is uni-modal (Fig. 3C) with a mean at 743±92 nm (mean±STD). While the distance between this transgene and *eve* fluctuates during the 30 min interval, there is no instance in which sustained physical proximity is established.

When we further tested the effect of reversing the orientation of *homie* in the original transgene (Fujioka et al., 2016), such that the *lacZ* reporter is downstream of *homie*, we find that pairing still occurs, but the regulation of the reporter by the *eve* enhancers is disrupted. We analyzed movies of 761 nuclei across 10 embryos for this *parS-homieR-lacZ* transgene. As expected, the corresponding bi-modal distribution resembles that for the regular *homie* transgene and has two means at 739±104 nm and 334±63 nm(mean±STD), respectively (Fig. 3D).

We next scored the nuclei with respect to the transcriptional activity of the *lacZ* reporter. Strikingly, all of the *parS-homie-lacZ* transgene nuclei showing *lacZ* transcription (i.e. presence of red signal, N=192) have a physical separation that falls within the distribution corresponding to the bound (i.e. homie-linked) conformation (Fig. 3A and 3B). But there are no nuclei in the unbound conformation that also express *lacZ*. Taken together, the above results demonstrate that *homie* pairing creates a local chromatin conformation that is permissive to transcription events by ensuring physical proximity between the *eve* enhancers and the promoter of *lacZ.* Consistent with this view, none of the nuclei in control *parS-lambda-lacZ* embryos express *lacZ.* From these observations we conclude that the *eve* enhancers must be in close proximity to the *lacZ* promoter in order to activate transcription.

**Fig 3:**
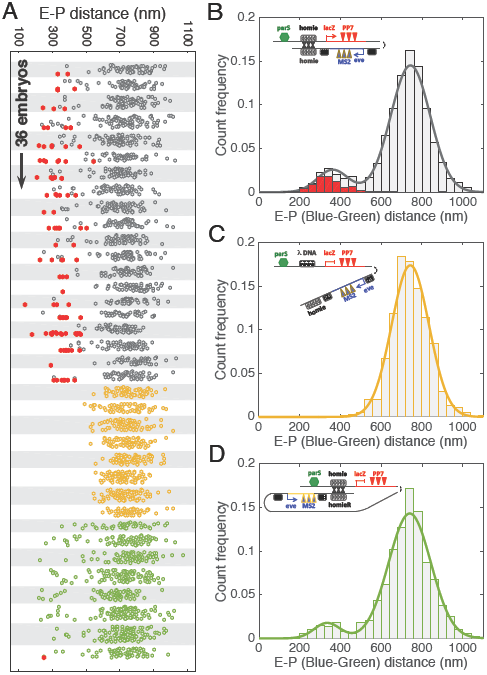
Physical proximity between enhancers and promoter is required to activate transcription. **(A)** For 36 embryos during nuclear cycle 14 (arranged vertically) the distribution of the time-averaged root-mean-squared (RMS) distance (E-P distance) between blue MS2 and green *parS* DNA foci is depicted in vertical scatter plots. Each data point corresponds to an individual nucleus displaying *eve* activity in embryos expressing *parS-homie-lacZ* (gray), *parS-lambda-lacZ* (orange) and *parS-homieR-lacZ* (green), respectively. Nuclei displaying *lacZ* activity are marked in red. **(B-D)** Distributions of RMS distances for three constructs: *parS-homie-lacZ* **(B)**, *parS-lambda-lacZ* **(C)**, *parS-homieR-lacZ* **(D)** with Gaussian fits. Insets show the genomic constructs in their presumptive conformations. See also Fig. S3.

### Necessity for sustained physical association

To assess the temporal relationship between enhancer–promoter proximity and transcriptional activation, we examine the dynamics of physical proximity and activation along our measurement window, in singe nuclei. All nuclei displaying a switch from OFF-to-ON (N=65) were aligned with respect to the time point when nascent transcripts could first be detected (Fig. 4A, see Methods). The data show a sharp switch in activity state with similar rates to those previously reported for nuclei exiting mitosis (Garcia et al., 2013). For the same data set, we then measured the mean distance between the green *parS* tag in the *parS-Homie-lacZ* transgene and the *eve* gene (blue) as a function of time by averaging across all OFF-to-ON time courses with the same alignment (Fig. 4B). We observed a continuous spatial convergence until the onset of transcription at which point the mean distance corresponds to an average separation of about 340 nm. These findings indicate that there is a close connection between the establishment of enhancer–promoter proximity and enhancer activation of transcription.

We also assessed the temporal relationship between the ON-to-OFF switch in transcription and the physical distance between the enhancer–promoter pair. In 42 nuclei that switch their transcriptional activity OFF during our observation window, a drop in transcriptional activity of the *lacZ* reporter is accompanied by an increase in the mean distance between the *parS-Homie-lacZ* transgene and the *eve* gene (Figs. 4C and 4D). Our experiments indicate that there is about a 4 min gap between the time when the *parS-Homie-lacZ* transgene first begins to detectably separate from the *eve* gene and a clear decline in transcriptional activity. Since our reporter gene is 5.5 kb in length and the measured RNA polymerase II (Pol II) elongation rate is 1.50.1 kb/min (Garcia et al., 2013), the major part of this delay must be due to engaged polymerases that continue to elongate after the *parS-Homie-lacZ* becomes unlinked from the *eve* enhancers. These results fit with a model in which sustained physical association of *eve* enhancers and the *lacZ* promoter is necessary for continuous initiation of transcription.

### Physical enhancer–promoter engagement leads to distinct topological conformations

While we have demonstrated that enhancer action depends upon sustained physical proximity to the promoter of the *lacZ* reporter, the proximity generated by *homie* pairing alone is not sufficient. Among the 307 *parS-homie-lacZ* nuclei in which *homie*-pairing likely occurred (i.e. nuclei occupying the short-distance peak in the histogram in Fig. 3B), only 63% express *lacZ*-PP7 during the 30 min window of observation in n.c. 14. Notably, in the nuclei that show reporter activity the MS2-parS distance is 27±8 nm (mean±SEM) lower than the distance in the nuclei where the *lacZ* reporter is inactive, suggesting that an additional compaction is associated with reporter activation.

The insufficiency of close proximity is even clearer when we employ the *parS-homieR-lacZ* reporter. This reporter still facilitates homie-pairing, yet likely renders the *lacZ* and the *eve* enhancers on opposite sides of these paired boundary elements (Fig. 3D, Fujioka et al., 2016). Of the 54 nuclei in which the *lacZ* transgene is close to the *eve* locus, there are only three in which the *eve* enhancers activate reporter transcription, and only for brief periods of 3-5 min.

Is transcriptional activation associated with an additional step that promotes physical enhancer–promoter engagement? We have shown that *homie* pairing establishes a new topological conformation in which the *parS-homie-lacZ* transgene and the *eve* locus are in close proximity (Fig. 3B). However, because the *eve* enhancers are distributed within 20kb of the eve-MS2 locus (Fig. 5A), *homie* pairing does not necessarily yield a close proximity between *eve* enhancers and the *lacZ* reporter. To further examine this point we take advantage of the property that independent *eve* enhancers regulate individual stripes of the *eve* pattern along the embryo (Fig. 5A). Thus by examining nuclei from different stripes separately we are able to explore the topology of the locus under different activating enhancers.

**Fig 4:**
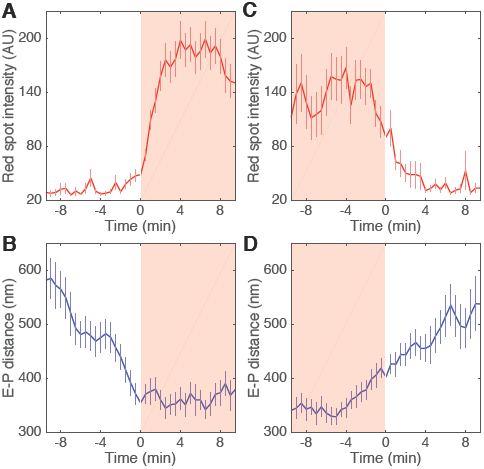
Dynamics of chromatin movement underlies kinetics of enhancer–promoter interactions and transcriptional activation. **(A)** Average *lacZ* activity as a function of time for nuclei turning on (time traces for individual nuclei are aligned such that activity starts at 0 min, i.e. first occurrence of red signal). **(B)** RMS distance between blue (MS2) and green (parS) DNA foci (E-P distance) in nuclei corresponding to data in (A). **(C)** Average *lacZ* activity for nuclei turning off, aligned such that activity ends at 0 min. **(D)** RMS distance between blue (MS2) and green (parS) DNA foci in nuclei corresponding to data in (C). In all panels shaded background signifies presence of *lacZ* activity.

We first considered the distance of the eve-MS2 gene relative to the *parS* tag (blue-green) in nuclei where *homie* pairing occurs but *lacZ* transcription is lacking (Red-OFF, Fig. 5B). We observed different distances in nuclei belonging to different stripes, i.e. a dependence of the spatial arrangement on the identity of the activating enhancers (Fig. 5C, Red-OFF). If an activating enhancer is physically engaged with the endogenous *eve* promoter, it co-localizes with the blue eve-MS2 focus. Thus the distance to the *homie* pair (green *parS* tag) should depend on the distance between the activating enhancer and the endogenous *homie*. Indeed, the distance between the *eve* gene and the *parS* tag of the inactive *lacZ* reporter in stripe 5 is shorter than that observed for nuclei in stripes 4/6 and 3/7, for which the enhancers are located farther away from the *parS* tag (Fig. 5C). Moreover, for the two enhancers that drive *eve* expression in a pair of stripes, 4/6 and 3/7, the distance between *eve* and the *parS* tag is closely matched within each stripe pair. Finally, for embryos with the *parS-homieR-lacZ* transgene the same trend is observed in nuclei with *homie* pairing (Fig. S4A). Taken together, these results argue strongly that the *eve* enhancers directly engage the endogenous *eve* promoter to activate transcription and that in each *eve* stripe a distinct topological conformation is adopted.

**Fig 5:**
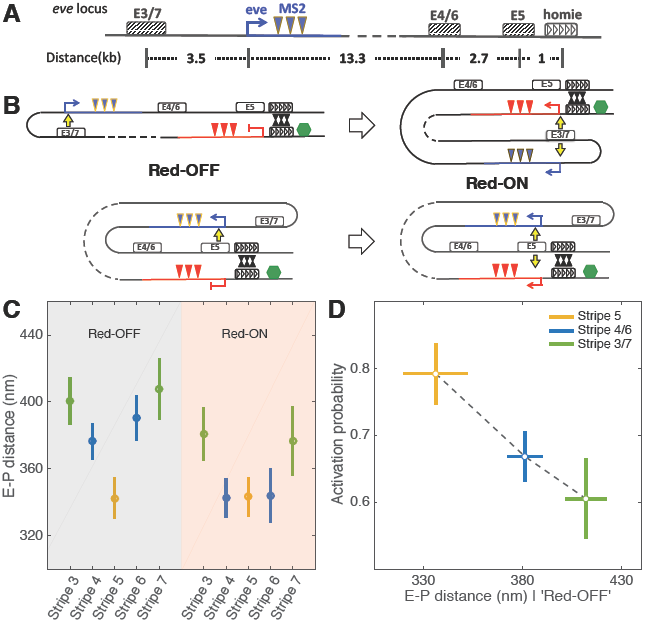
Activation from endogenous enhancers is governed by enhancer–promoter distances. **(A)** Endogenous *eve* locus with stripe-3/7 enhancer, stripe-4/6 enhancer and stripe-5 enhancer and their relative locations with respect to the *eve* promoter and the endogenous homie sequence. The stripe 5 and stripe 4/6 enhancers are located downstream of the *eve* promoter; 1.0 kb and 3.7 kb upstream of, *homie*, respectively. The stripe 3/7 enhancer is located upstream of the *eve* promoter; 20.5 kb from homie. **(B)** Cartoon showing *eve* locus under different looping conformations (Red-OFF and Red-ON) after *homie-homie* pairing. Top row is for stripe 3/7; bottom row is for stripe 5. **(C)** RMS distance between blue (MS2) and green (parS) DNA foci (E-P distance) for all nuclei in which *homie-homie* pairing occurs. RMS distances are calculated for individual *eve* stripes. Red-OFF and Red-ON correspond to the *eve* locus conformations illustrated in (B) for the *parS-homie-lacZ* transgene. **(D)** Fraction of *homie*-linked nuclei in each stripe that express *lacZ* (activation probability) is plotted as a function of distance as measured in (C). See also Fig. S4.

Next, if transcriptional activation entails physical engagement, in each stripe we expect that in nuclei where *lacZ* transcription is active (Red-ON, Fig. 5B) not only will the eve-MS2 gene be in close proximity to the activating enhancer but also the *parS-homie-lacZ* reporter. Indeed, in the nuclei with the active reporter a significant shortening of the distance between the *eve* enhancers and the *lacZ* promoter is observed in stripes 4/6 and 3/7 when *lacZ* is activated (Fig. 5C), which further argues for transcription associated compaction of the locus. Critically, there is no *lacZ* activity dependent shift apparent for stripe 5 nuclei as in these nuclei the *eve* stripe 5 enhancer is already in close proximity to the *lacZ* promoter.

Finally, a stripe-specific topology is also favored by observations in measurements of the activation probability of the promoter driving *lacZ* expression. If transcription of the homie-linked reporters is linked to enhancer engagement, a plausible expectation is that activation frequency would be distance dependent. To assess the effect of distance we determined the fraction of *homie*-linked nuclei in each stripe that express *lacZ.* As predicted, the fraction of transcriptionally active reporters decreases with increasing distance between the stripe enhancer and the *lacZ* promoter (Fig. 5D). Stripe 5 has the highest activation probability ( 80%), while stripes 3/7 have the lowest probability of enhancer engagement with the *lacZ* reporter. The fact that we observe a linearly decaying relationship is consistent with contact probability measurements (Lieberman-Aiden et al., 2009). Together, these results provide compelling evidence that transcriptional activation entails physical enhancer–promoter engagement and is thereby associated with stripe-specific topological compaction of the *eve* locus.

### Promoter competition has phenotypic consequences

In our experiments, the *eve* stripe enhancer drives expression from two different *eve* promoters, one for the endogenous *eve* gene and the other for the *lacZ* reporter (Fig. 5B, Red-ON). If the activity of the stripe enhancers is limiting, the *lacZ* reporter will reduce transcription of the *eve* gene. To determine whether promoter competition occurs in our genomic setup we compared *eve* transcription (i.e. the intensity of the blue MS2 signal) in individual nuclei in which *lacZ* is active and nuclei in which *lacZ* is silent (Fig. 6A, see Methods). For each *eve* stripe, we measured a 5%-25% reduction in endogenous *eve* transcription in nuclei in which *lacZ* is also transcribed compared to nuclei in which *lacZ* is not transcribed. The average reduction per nucleus is highest for stripe 5, and lowest for stripes 3 and 7.

*Eve* is a primary pair-rule gene that is responsible for segment patterning. To determine whether the observed reduction in *eve* transcription has any phenotypic consequences, we crossed males carrying a tagless *homie-lacZ* transgene at −142 kb (Fig. 1A) to females heterozygous for a wild type *eve* gene and an *eve* deficiency *(Df(2R)eve*). *eve* is weakly haploinsufficient and 7% of *+/Df(2R)eve* flies display patterning defects in even-numbered abdominal segments. Consistent with the reduction in the level of *eve* nascent transcripts, the presence of the *homie-lacZ* transgene exacerbates *eve* haploinsufficiency (Fig. 6B-D). Altogether 29% of the *Homie-lacZ/Df(2R)eve* flies have abdominal defects which corresponds to a 4-fold increase (Fig. 6E). Taken together, the above results suggest that competition between two promoters at the transcriptional level in the early embryo has phenotypic consequences for patterning in the adult. These findings provide evidence that manipulating topological chromatin structures can interfere with developmental programs.

## III. DISCUSSION

Despite extensive studies over more than three decades, many questions still remain regarding how enhancers communicate with their target promoters over large genomic distances. Critically, although recent FISH and 3C-based genomic experiments have provided extensive evidence supporting physical interactions for long-range enhancer–promoter communication (Deng et al., 2012; Fabre et al., 2015; Hnisz et al., 2016; Ji et al., 2016; Phillips-Cremins et al., 2013; Rao et al., 2014; Williamson et al., 2016), we still miss a dynamic picture that could disentangle cause from consequence and that could distinguish transient contact with formation of stable topological structures. Here, we have developed a live imaging approach on a well-studied endogenous locus that allows us to probe the dynamics of topological chromatin structures at the single-cell level and its impact on transcriptional activity. We show directly that sustained physical proximity is necessary for enhancer action, and distinct topologies are observed for different pairing of the promoter with an activating enhancer, even when enhancers are clustered together. Though all our results are based on a specific experimental system it seems likely that they will be more broadly applicable.

**Fig 6:**
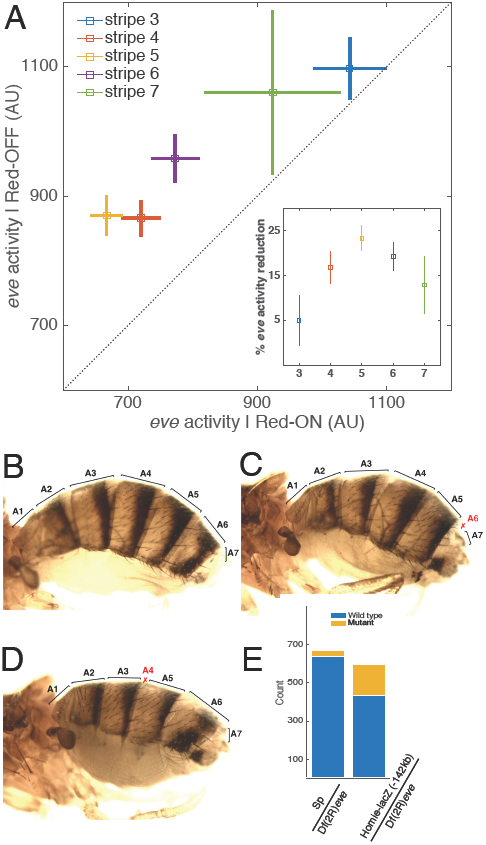
Long-distance-mediated promoter competition results in patterning phenotypes. **(A)** Endogenous *eve* activity in nuclei that also display *lacZ* activity (x-axis) is lower than in nuclei where *lacZ* is not expressed (y-axis). Inset: Reduction in *eve* activity for each stripe. **(B-D)** Adult wild-type (B) and mutant (C and D) flies from crosses between Sp/*Homie-lacZ* males and CyO/Df(2R) *eve* females. (C) and (D) show defects in abdominal segments A4 and A6, respectively, resulting from reduced *eve* activity in stripe 5 and stripe 6, respectively. Abdominal segments are labeled, with defective segments marked in red. **(E)** Results of phenotype scoring. Mutant counts include both A4 and A6 phenotype. p value from Fishers exact test.

To examine long-range transcriptional activation we placed a reporter gene at 150 kb distance of the presumptive *eve* enhancers. Although such distance is unusually large for enhancer–promoter interactions in flies, in higher eukaryotes enhancers are known to function over comparable or greater distances (Sanyal et al., 2012). At such distance the chromatin fiber can display fast random movements, which creates an entropic difficulty for specific long-range chromatin interactions and thus a kinetic barrier for the establishment of a productive pre-initiation complex. Architectural proteins have been suggested to increase contact frequency between chromatin loci separated by large genomic distances and facilitated the formation of sustained physical proximity permitting functional enhancer–promoter interactions (Erokhin et al., 2011; Shen et al., 2012; Symmons et al., 2014). The subdivision of genomes to topologically associating domains (TADs) was further suggested to facilitate local (intra-domain) chromatin interactions and reduce the search space, as exemplified for VDJ recombination in the mammalian immune system (Lucas et al., 2014).

To facilitate the interaction between the enhancers and the reporter in our experimental setup, we thus employ the *homie* element in our transgene, thereby inducing a stable loop conformation with the endogenous *homie* element that demarcates the 3 end of the *eve* locus. This element was shown to be bound by a wide array of Drosophila insulator proteins in the blastoderm embryo, including Su(Hw), CP190, BEAF-32 and dCTCF (Negre et al., 2010). homie-*homie* pairing between the transgene and the *eve* locus provides spatial proximity that we find to be necessary to allow the enhancers to activate transcription of the reporter. When the *homie* element in the transgene is replaced by lambda DNA, transcriptional activity of the reporter gene is not observed. As this transgene should encounter the *eve* enhancers with roughly the same frequency as the transgenes carrying homie, it demonstrates that transient encounters are not sufficient to activate PolII transcription, and the various transcription factors associated with the *eve* enhancers and the *eve* promoter are unable to establish stable and/or productive enhancer–promoter contacts on their own.

Another example in which an architectural element is deployed de novo to stabilize long distance enhancer–promoter interactions is the variably occupied tissue specific dCTCF binding site in the Ultrabithorax intron just upstream of the abx/bx enhancer (Magbanua et al., 2015). More generally, TAD boundaries in mammalian genomes often harbor CTCF binding sites, which are believed to play a major role in genome organization. Intriguingly, these sites were recently shown to have a specific orientation (Guo et al., 2015; Rao et al., 2014). When we examine a reporter construct in which we inverted *homie* orientation we find that while pairing is still maintained, the frequency of transcriptional activation drops significantly. The cause is likely because the enhancers and the promoters are now found on opposite sides of the paired insulators (Fujioka et al., 2016). Thus, alteration to these binding sites can bare important functional implications and indeed many cancer types accumulate CTCF binding site mutations that modulate chromatin conformations with subsequent influences on transcription (Flavahan et al., 2016; Katainen et al., 2015). Our experimental setup provides an opportunity for exploring the dynamic relationship between topological alterations and transcriptional regulation resulting from mutations in boundary elements.

Overall, our experiments demonstrate that physical proximity is a necessary condition for the *eve* enhancers to activate *lacZ* expression. Importantly, we find that transcription ceases upon dissociation of the enhancers and the reporter, suggesting a requirement for sustained physical proximity, and excluding mechanisms of transient enhancer promoter associations leading to stable activity. While we observe this tight relationship between physical proximity and activation, for other genes, loop formation could be a permissive step, with pre-formed enhancer–promoter interactions preceding transcriptional activation (de Laat and Duboule, 2013; Ghavi-Helm et al., 2014; Jin et al., 2013; Montavon et al., 2011). This discrepancy might reflect regulatory strategies that are gene or developmental context dependent. Nevertheless, consistent with the observations that a large amount of reorganization in enhancer–promoter interactions exist upon tissue differentiation (Dixon et al., 2015; Javierre et al., 2016), our results show that establishment of physical interactions between enhancers and promoters could be a rate-limiting step for transcriptional activation and play a key role in regulating transcription and cell fate.

Applying our single-cell live imaging approach in the early fly embryo offers an unprecedented opportunity to examine topological chromatin structures of a locus in a cell-specific way. We provide compelling evidence that *eve* expressing cells adopt distinct topological chromatin structures according to the stripe-specific identity of the activating enhancer. Notably the *eve* enhancers are located within 10 kb of the endogenous *eve* promoter and the *homie*-paired promoter of the reporter gene. A possible model could be that the adjacent *eve* enhancers will be found in close proximity to the promoters within a local environment permitting any of the enhancers to activate either of the promoters without further topological distinctions. Our observation of a stripe-specific distance between the locus of the *eve* gene and the *homie*-tethered *parS* tag, and the further compaction of the locus upon concurrent activation of the reporter, seem to exclude such a model. But rather this seems to suggest the necessity of a specific enhancer–promoter engagement for transcriptional activation, even within a relatively short genomic range.

What is the spatial scale that defines enhancer–promoter engagement? Transcriptional activity in our experimental system is only observed when the reporter moves into close proximity to the *eve* locus. The transition from inactivate to active occurs when the parS tag comes within 350 nm of the *eve* gene. When the reporter moves more than 350 nm away from the *eve* gene, transcription shuts off, and already engaged Pol II bring their elongation process to completion. The range of the observed necessary physical proximity is likely to be dictated by the spatial scale set up by the size of the protein complexes that are believed to physically bridge enhancers with promoters (¿2MDa with diameter ¿20nm (Dill et al., 2011)), for example Mediator complex and the Pol II pre-initiation complex (Kagey et al., 2010; Plaschka et al., 2015; Robinson et al., 2016). This requirement of 350 nm in enhancer–promoter proximity in order to activate transcription is comparable with DNA FISH measurements in mammalian systems that estimate the proximity between the shh promoter and its enhancers in mouse limbs (Amano et al., 2009); (Williamson et al., 2016).

Finally, when the endogenous *eve* locus is presented with an extra ectopic promoter, its activation results in concomitant reduction in endogenous *eve* transcription, phenotypically leading to patterning defects in the corresponding segments. Usually, competing promoters are physically linked within short genomic distances (¡20kb) (Foley and Engel, 1992; Fukaya et al., 2016). However, our results show that physically associating the competing promoters could in principle also be achieved through dynamic alterations of chromatin conformations. This observation reinforces the notion that rearrangement of topological chromatin structures is able to re-wire enhancer–promoter interactions, and that the rewiring can cause new phenotypes in many developmental contexts. For example in humans, structural chromosome variants that disrupt boundary elements flanking TADs may lead to diseases (Franke et al., 2016; Lupianez et al., 2015). Thus our experiments provide a potential path to synthetically rewire enhancer–promoter interactions to perturb chromatin topologies and thereby interfere with developmental programs (Deng et al., 2012; Deng et al., 2014),

In conclusion, we propose a methodological framework for investigating the relationship between chromatin dynamics and long-range enhancer-mediated transcriptional regulation. With the ability of tracing endogenous loci in individual cells and simultaneously measuring temporal dynamics and spatial information, new mechanistic insights into enhancer–promoter interactions are likely to be uncovered.

### Methods Summary

Plasmid construction

Transgenic fly generation

Fluorescent in-situ hybridization

Phenotypic scoring

Microscopy and imaging conditions

Image processing and data analysis

## Acknowledgements

We thank P. Schedl for bringing the *eve-homie* system to our attention. We thank K. Bystricky for introducing us to the ParB/ParS system, and F. Payre and P. Valenti for sharing a ParB-eGFP plasmid and the *parS* sequence. We thank P. Ratchasanmuang and S. Ryabichko for assistance with cloning and fly husbandry, and the Bloomington Drosophila Stock Center for fly strains. We are also grateful to L. Barinov, S. Blythe, M. Levine, M. Levo, P. Schedl, E.F. Wieschaus for discussion and help with the manuscript. This study was funded by grants from the National Institutes of Health (U01 EB021239, R01 GM097275, and R01 GM050231). HC was supported by the Charles H. Revson Biomedical Science Fellowship.

---

**SUPP. FIG. 1:**
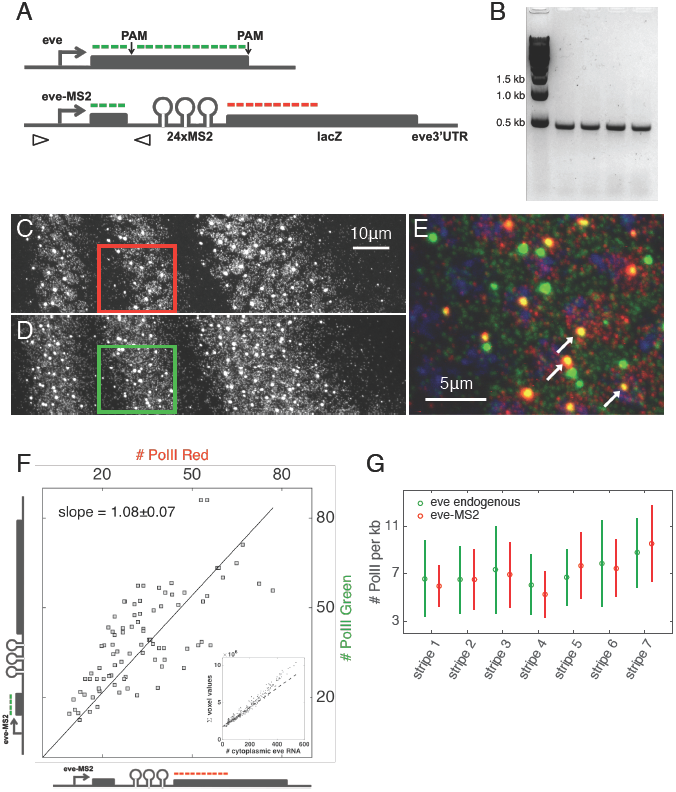
*eve-MS2* allele recapitulates the expression pattern and transcriptional activity of the endogenous *eve* gene. **(A)** Editing the endogenous *eve* locus (upper) to obtain the *eve-MS2* allele (bottom). Arrowheads indicate primers for PCR genotyping. Green and red lines mark sequences targeted by smFISH probes (eve-atto633 and lacZ-atto565, respectively). (B) Genotyping the *eve-MS2* allele. PCR results from four independent transformant lines are shown. **(C-G)** FISH quantification of the transcriptional activity of the *eve-MS2* allele. Maximum Z-projections are shown for lacZ-atto565 channel (C) and eve-atto633 channel (D) of an *eve-MS2/eve+* embryo. Three *eve* stripes (stripe 5-7 from left to right) at 45 min in nuclear cycle 14 are shown. (E) Magnified view of square in C/D. Arrows indicate *eve-MS2* transcription loci that are labeled by both probes. (F) Cytoplasmic spots and active transcription spots are identified by image analysis routines (see Methods). A cytoplasmic unit (CU) that corresponds to fluorescent intensity of a single cytoplasmic mRNA is extracted. Panel shows number of RNA polymerase II (Pol II) on the *eve-MS2* loci as in inferred from either the CU derived from lacZ-atto565 (x-axis) or from eve-atto633 (y-axis) measurements. Inset shows calculation of cytoplasmic unit for *eve.* Specifically a sliding window of 220x220x23 pixel (16.5x16.5x7.4 *μm*^3^) was applied to the raw image stack (C and D) and the total pixel values in the window were plotted against the number of cytoplasmic spots found in the window. A linear fit in the range of 0-100 cytoplasmic spots was applied to extract CU for each probe set. (G) Comparison of the PolII number on the *eve-MS2* locus and on the endogenous *eve* locus. Note that the numbers reported in (F) and (G) are for two sister chromatids.

**SUPP. FIG. 2:**
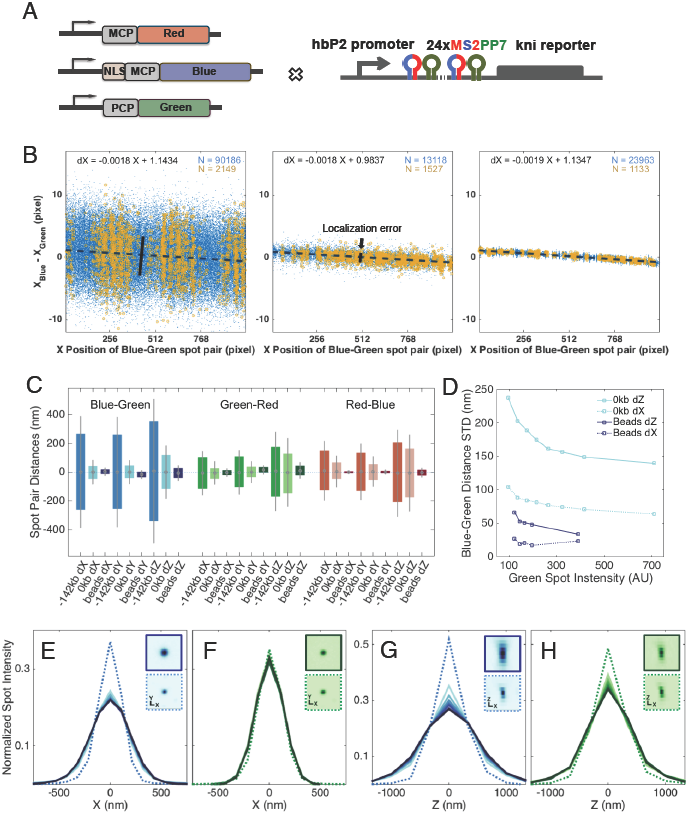
Spot localization precision and measurement error. **(A)** Genetic design of a transgene that co-localizes all three reporter systems. MS2 and PP7 stem loops are alternated and repeated 24 times. A *kni* reporter gene is driven by a *hunchback* P2 (hbP2) promoter, resulting in expression in all nuclei located in the anterior 10-45% of the embryo. **(B)** A sample data set accompanying the methods for chromatic aberration correction and measurement error determination. Panels show the linear distance (along x-coordinate only) for each blue-green spot pair as a function of the pairs x-position for embryos carrying the *parS-homie-lacZ* transgene (left), embryos carrying the three-color co-localization transgene from (A) (middle), and TetraSpec beads (right), respectively. Blue data points are for all spot pairs at all time frames for all embryos analyzed. Yellow data points are from one of the embryos (or one set of experiment for the beads). Linear fits in each panel report on the chromatic aberrations between blue and green spots in the x-direction. As slopes and intercepts for the different samples show no significant differences, chromatic aberrations can be corrected for each individual embryo data set internally. **(C)** Summary of the distributions of spot pair distances (after chromatic aberration correction) for the three configurations in (B). Each direction (x, y, and z) is shown for each color combination. For example, for the blue-green (MS2-parS) distances on the x-direction the STD of the *parS-homie-lacZ* transgene (labeled −142kb) corresponds to the solid black bar shown in the left panel of (B). Spot localization errors are estimated from the STDs measured with the three-color co-localization control embryos (labeled 0kb). Solid lines: standard deviation (STD); bars: 25%-75% quantiles. **(D)** Dependence of localization precision on signal intensities. Since localization precision scales with the square root of the number of photons, we directly compare localization errors from the three-color co-localization control embryos with the errors measured from immobilized beads of the same fluorescent intensity values. Panel shows that the differences in the localization errors between embryos and beads are not due to difference in photon counts, and thus about 2/3 of that localization error is due to motion blurring of the moving spot during acquisition. The remaining 1/3 (i.e. error obtained from immobile beads) stems from optical measurement noise and the analysis pipeline. **(E-H)** Optical characterization of transcription and parS spots. For each fluorescent channel, all identified spots are classified into eight groups according to their intensities. An average spot for each group is created by aligning all spots so that the brightest pixels are at the center of a 25x25x13 voxel region of interest (ROI) and taking the average intensity per voxel in that region over all spots. The intensity profiles along the X- (E and F) and Z-cross-sections (G and H) for the blue MS2 average spot (E and G) and green parS average spot (F and H) are plotted (darker curves represent brighter spots). Dashed lines are from equivalent measurements of TetraSpec beads. Images of the average spots for the brightest blue (or green) MS2 spots (upper) and for the beads (lower) are shown as panel insets.

**SUPP. FIG. 3:**
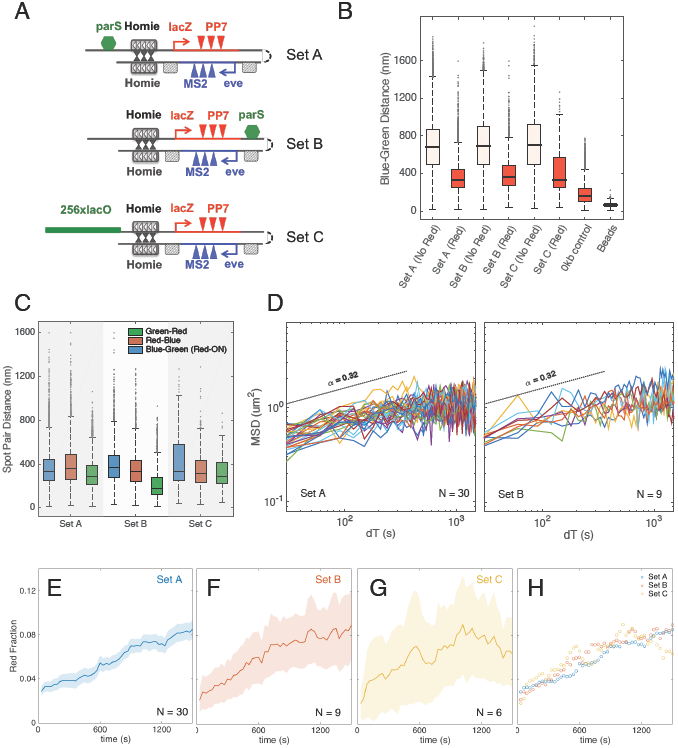
Different genomic labeling approaches report on similar chromatin dynamics and transcription kinetics. **(A)** Three methods of labeling genomic loci. **(B)** The measured Blue-Green (MS2-parS) distances are not sensitive to labeling approaches. Boxplot showing distributions of the instantaneous distance between spot pairs in the same nuclei. Distances shown are after chromatic aberration corrections. For all three genomic settings, the MS2-parS (blue-green) distances show no significant differences. This is observed regardless the absence (No Red) or presence (Red) of *lacZ* activity. The whiskers extend to 1.5 times the inter quantile range from the 25% and the 75% quantiles. **(C)** The distances between spot pairs reflect their genomic arrangement. Distributions of the instantaneous distance between spot pairs are plotted. Box and whiskers as in (B). Distances shown are after chromatic aberration corrections. Notice that the *parS*-PP7 (green-red) distance is significantly shorter when the *parS* tag is located at the 3 side of the *lacZ* repoter. **(D)** Mean square displacement (MSD) plots for Set A and Set B. Each MSD trace is a result of population ensemble of all nuclei in a single embryo (embryo-averaged MSD, see Methods). Results from the two genomic settings display sub-diffusive characteristics with a scaling power of 0.32 and their anomalous diffusion coefficients show no significant difference. **(E-H)** Transcriptional activation of *lacZ*-PP7 is not affected by labeling approaches. The fraction of *eve*-expressing nuclei that also contain active *lacZ*-PP7 is plotted as a function of time for three genomic settings. Apparently, the presence or locations of *parS* tags do not interfere with enhancer actions and transcriptional activation.

**SUPP. FIG. 4:**
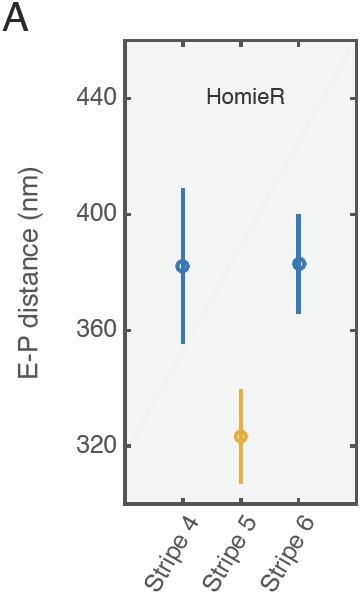
Stripe-specific topological difference for the *parS-homieR-lacZ* transgene. **(A)** RMS distance between blue (MS2) and green (parS) DNA foci (E-P distance) for all nuclei in which *homie* pairs with the reversed *homie* (homieR) sequence. RMS distances are calculated for individual *eve* stripes.

